# *CBP/p300* orthologs *CBP2* and *CBP3* play distinct roles in planarian stem cell function

**DOI:** 10.1101/2020.09.08.287417

**Authors:** Clara R. Stelman, Britessia M. Smith, Bidushi Chandra, Rachel H. Roberts-Galbraith

## Abstract

Chromatin modifications function as critical regulators of gene expression and cellular identity, especially in the regulation and maintenance of the pluripotent state. However, many studies of chromatin modification in stem cells—and pluripotent stem cells in particular—are performed in mammalian stem cell culture, an *in vitro* condition mimicking a very transient state during mammalian development. Thus, new models for study of pluripotent stem cells *in vivo* could be helpful for understanding the roles of chromatin modification, for confirming prior *in vitro* studies, and for exploring evolution of the pluripotent state. The freshwater flatworm, *Schmidtea mediterranea*, is an excellent model for studying adult pluripotent stem cells, particularly in the context of robust, whole-body regeneration. To identify chromatin modifying and remodeling enzymes critical for planarian regeneration and stem cell maintenance, we took a candidate approach and screened planarian homologs of 26 genes known to regulate chromatin biology in other organisms. Through our study, we identified six genes with novel functions in planarian homeostasis, regeneration, and behavior. We also identified in our list five planarian homologs of the mammalian CREB-Binding Protein (CBP) family of histone acetyltransferases, representing an expansion of this family in planarians. We find that two planarian CBP family members are required for planarian survival, with knockdown of *Smed-CBP2* and *Smed-CBP3* causing distinct defects in stem cell maintenance or function. Loss of CBP2 causes a quick, dramatic loss of stem cells, while knockdown of CBP3 more narrowly affects stem cells, preferentially decreasing markers of neural progenitors. We propose that the division of labor among a diversified CBP family in planarians presents an opportunity to dissect specific functions of a broadly important histone acetyltransferase family in stem cell biology.

## Introduction

Stem cell maintenance and differentiation require coordinated gene expression, which can be achieved by changing chromatin structure through covalent posttranslational histone modifications and nucleosome remodeling (For review, see: (Adam and Fuchs, 2016; Di Giammartino and Apostolou, 2016; Yadav et al., 2018). Histone modifications include, but are not limited to: methylation, acetylation, phosphorylation, and ubiquitination (Kouzarides, 2007). Each modification occurs in specific combinations on histone tails to modulate local gene regulation (Strahl and Allis, 2000). In addition to histone modifications, histone remodeling enzymes physically rearrange nucleosomes and facilitate removal or replacement of histone proteins (Narlikar et al., 2013). Stem cells, particularly pluripotent stem cells, possess a unique epigenetic landscape that allows them to maintain chromatin poised for acquisition of specific fates (Bernstein et al., 2006; Gafni et al., 2013; Mikkelsen et al., 2007; Pan et al., 2007; Shema et al., 2016). Thus, enzymes that modify and remodel chromatin play key roles in maintenance of and exit from the pluripotent state (e.g. (Ho et al., 2009; Onder et al., 2012)). However, it is difficult to study maintenance and exit from pluripotency *in vivo* in traditional model systems due to the early and transient presence of pluripotent cells in most organisms during embryogenesis.

Planarians offer an attractive model for studying the biological mechanisms underlying pluripotent stem cell maintenance and differentiation *in vivo*, due to the presence and persistence of pluripotent stem cells in the adult body. In response to amputation, planarian stem cells divide to form a blastema, an unpigmented tissue within which missing structures are generated (Baguñà et al., 1989). The identification of stem cell progeny markers (Eisenhoffer et al., 2008; Scimone et al., 2014; van Wolfswinkel et al., 2014) and the development of experimental techniques such as *in situ* hybridization (ISH; (Umesono et al., 1997)) and RNA interference (RNAi; (Sánchez Alvarado and Newmark, 1999)) provided an opportunity to identify and study genes important for planarian stem cell function. Studies have reported roles for diverse genes including *bruli* and *piwi* homologs in regulation of stem cell proliferation and self-renewal (Guo et al., 2006; Palakodeti et al., 2008; Reddien et al., 2005b), but the full complement of genes required for planarian stem cell function is still being identified.

So far, several planarian genes encoding chromatin remodeling enzymes or chromatin modifiers have been shown to regulate stem cell function in planarians (Dattani et al., 2019; Robb and Sanchez Alvarado, 2014). Knockdown of some histone methyltransferase genes from the SET1-MLL family causes reduced regeneration, specifically due to the roles these genes play in stem cell maintenance and mitosis (Duncan et al., 2015; Hubert et al., 2013). Members of SWI/SNF complexes are also important for planarian stem cell biology (Trost et al., 2018). In addition to functioning in stem cell maintenance, chromatin modifying and remodeling enzymes also fulfill key functions during differentiation of planarian stem cells. A Mi-2-like chromatin remodeling enzyme, encoded by *Smed*-*CHD4*, regulates the development and differentiation of stem cell progeny to maintain proper tissue turnover and regeneration (Scimone et al., 2010). Other factors, including p66 and RbAp48-like, also play roles in differentiation (Bonuccelli et al., 2010; Vásquez-Doorman and Petersen, 2016) and ciliogenesis (Duncan et al., 2015). More recently, planarian stem cells were also shown to possess bivalent chromatin marks at gene loci poised for expression during differentiation (Dattani et al., 2018). While chromatin-regulating proteins have been established to play important roles in planarian stem cell biology (Rossi et al., 2003; Zeng et al., 2013), nearly 100 conserved chromatin regulators are encoded by the planarian genome (Robb and Sanchez Alvarado, 2014) and many of these uncharacterized genes may play critical, unexplored roles in stem cells.

In this work, we took a candidate approach to identify potential chromatin regulators that function in planarian regeneration and/or stem cell biology. We identified two CREB-Binding Protein (CBP) family members and four other putative chromatin regulators that function in planarian homeostasis and regeneration. We find that the planarian CBP family is expanded and that two members, *CBP2* and *CBP3*, are necessary for stem cell viability and function. Specifically, we find that *CBP2* is required for stem cell maintenance whereas *CBP3* plays a relatively specific role, including an effect on neuronal progenitor markers. Our data reinforce the importance of the CBP family in stem cell biology and we propose that understanding the functional differences between planarian CBP genes may shed light on CBP family function in stem cells of other organisms.

## Methods

### Animal maintenance

Asexual planarians (CIW4) were maintained at 18-22°C in the dark. Animals were kept in Ziploc reusable containers in either 0.5g/L Instant Ocean Sea Salts (Spectrum Brands) dissolved in ultrapure deionized water or in Monjuïc salts as previously described (Cebrià and Newmark, 2005). Organic, pureed calf or beef liver was used to feed the animals once a week. Animals were starved for a minimum of 1 week prior to use in experiments.

### Identification of chromatin modifying genes and cloning

Planarian homologs of genes were identified using tblastn against the *Schmidtea mediterranea* genome (Robb et al., 2008). Full length segments were, when possible, determined using transcriptomic resources (Brandl et al., 2016; Kao et al., 2013). 500-750 bp segments of these genes were PCR-amplified from asexual cDNA using primers shown in Supplementary Table 1. Each PCR product was ligated into Eam1105I-digested pJC53.2 vector (Quick Ligation Kit, Roche) for use in ISH and RNAi experiments (Collins et al., 2010). Details on all genes from this study are listed in Supp. Table 1.

### CBP ortholog domain comparison and phylogenetic analysis

The Pfam 27.0 database (http://pfam.xfam.org/) was used to identify chromatin-acetylating Kat11 domains (Punta et al., 2012) in CBP proteins. The domains of CBP orthologs from *Drosophila melanogaster, Homo sapiens*, and *Schmidtea mediterranea* were compared. Phylogeny was analyzed using www.phylogeny.fr (Dereeper et al., 2008). The a la carte option was selected with MUSCLE for alignment (Edgar, 2004), Gblocks for curation (Castresana, 2000), and PhyML for construction of the phylogenetic tree (Guindon et al., 2010).

### RNAi Experiments

dsRNA was synthesized from clones described above via *in vitro* transcription as previously described (Chong et al., 2013; Rouhana et al., 2013), using primers in Supp. Table 2. Animals were fed 2-4 μg dsRNA mixed with 30-35 µl of 4:1 liver:salts mixture. These volumes were multiplied by 3 for larger experiments. Animals were kept in 60 mm (10-12 worms) or 100 mm (30-40 worms) Petri dishes, washed after feedings, and supplemented with 1:1000 gentamicin sulfate (50mg/ml stock, [Gemini Bio-Products]) throughout the experiment. dsRNA matching *green fluorescent protein* (GFP) or bacterial genes *ccdB* and *camR* was used for negative control treatments.

For our initial regeneration experiments (Fig. 1, 2B and Supp. Fig. 1), all but *CBP2*(RNAi) animals were fed on days 0, 6, and 12. These animals were cut pre-pharyngeally on day 17 and trunk pieces were allowed to regenerate for 5 days. On day 22, surviving animals were processed for immunofluorescence or ISH. For additional *CBP/CoREST(RNAi)* regeneration experiments (Fig. 2A, C) and *neuroD-1(RNAi)* experiments (Fig. 6F-I, Supp. Fig. 3), animals were fed on days 1, 7, and 13, amputated on day 19, and killed and fixed on day 25 (6 days post amputation -dpa). *CBP2(RNAi)* animals (Fig. 1, 5) were fed on days 0, 6, and 10 and were processed for RT-qPCR or ISH at the first sign of lysis (usually day 2-3 days after the final feeding). *CBP3(RNAi)* follow-up experiments were conducted as follows: animals in Fig. 6A-D, and 6J and Supp. Fig. 2 were fed on days 0, 6, and 10; for RT-qPCR (Fig. 6B), animals were amputated prepharyngeally on day 16 and processed on day 21; for ISH and blastema quantification (Fig. 6A, 6C-D, 6J and Supp. Fig. 2), animals were amputated prepharyngeally on day 17 and killed and fixed on day 23. For the survival curve experiment (Fig. 3A-C), animals were fed dsRNA once per week and observed twice per week for a period of 166 days.

**Figure 1.**
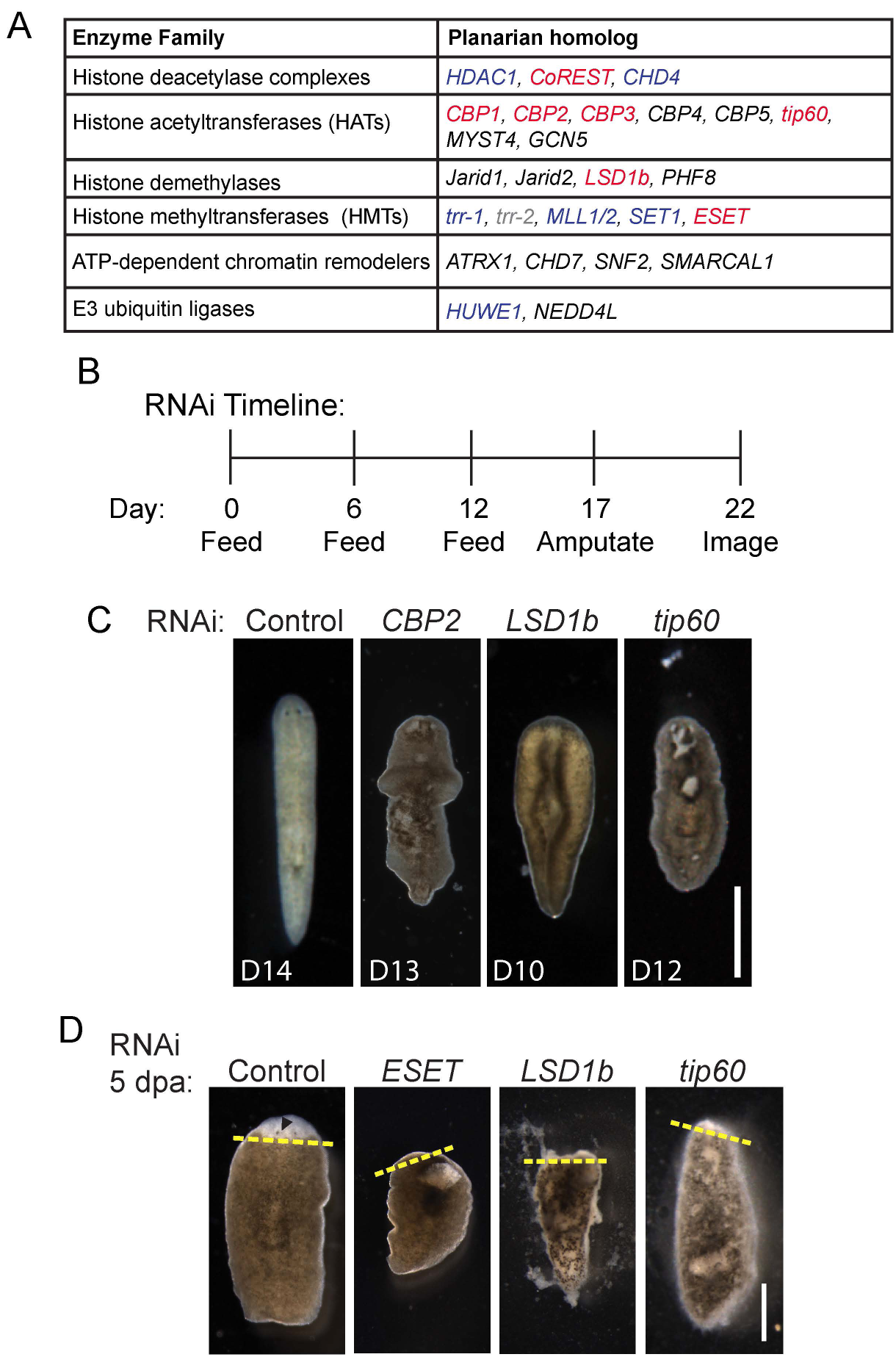
Chromatin modifying and remodeling enzymes are important in planarian homeostasis and regeneration. (A) A list of 26 planarian homologs of genes with roles in chromatin biology from our functional screen. Red indicates genes with RNAi phenotypes identified in this paper. Blue indicates previously published genes with phenotypes: HMTs (Hubert et al., 2013); *HDAC1* (Reddien et al., 2005a; Robb and Sanchez Alvarado, 2014; Zeng et al., 2013); *CHD4* (Scimone et al., 2010); and *HUWE1* (Henderson et al., 2015). Grey indicates previously published genes with no phenotypes: *trr-2* (Hubert et al., 2013). Black indicates genes identified in this paper with no overt RNAi phenotype (data not shown). (B) An RNAi time course (in days) for regeneration experiments performed in C (without amputation) and D (with amputation). (C) Prior to scheduled amputation, systematic lysis and/or curling was observed for *CBP2(RNAi), LSD1b(RNAi)*, and *tip60(RNAi)* animals. Representative images are shown. (D) Analysis of regenerating blastemas. Compared to control, RNAi animals developed either no or small blastemas, but lysis was also present for *ESET(RNAi), LSD1b(RNAi)* and *tip60(RNAi)* animals. Representative images are shown from 5 days post amputation (dpa). Dotted yellow lines represent blastema boundary. n=10 for each. Scale bar: 1 mm (C), 500 µm (D).

**Figure 2.**
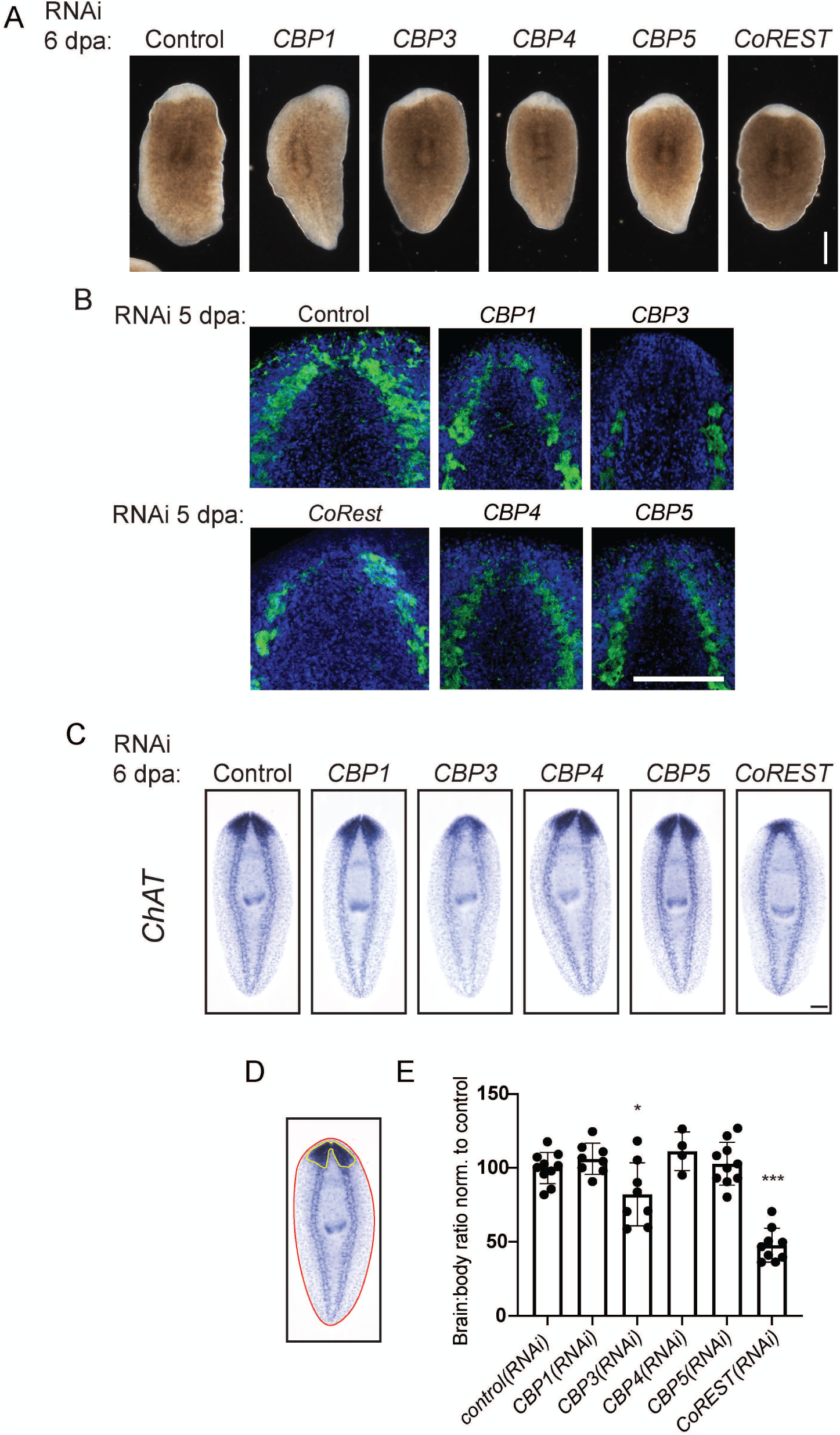
CBP3 and CoREST are required for proper regeneration. (A) Animals were treated with RNAi targeting the indicated genes. Blastemas formed at 6 dpa in all conditions, though they were slightly smaller after *CBP3(RNAi)* and *CoREST(RNAi)*. (B) After RNAi as in Fig. 1B, immunofluorescence (IF) staining was used to test the ability of animals to regenerate their nervous system after short-term RNAi. Regenerated animals were stained using an anti-synapsin antibody (green) that marks the animal’s neuropil and DAPI to mark cell nuclei (blue). *CBP3*(RNAi) animals regenerated cephalic ganglia with decreased anti-synapsin staining. (C) In order to quantify brain regeneration, RNAi-treated animals at 6 dpa were subjected to ISH with riboprobes matching *choline acetyltransferase* (*ChAT*). Smaller brains were observed after *CBP3(RNAi)* and *CoREST(RNAi)*. (D) To quantify brain size, we measured the area of the brain (yellow line) and body (red) in ImageJ to create a brain to body ratio (Roberts-Galbraith et al., 2016). This ratio was normalized so that *control(RNAi)* brain size was set to 100%. (E) Measurements of brain size as in (D) were conducted in triplicate for each animal and averaged across 4-10 animals per condition. There was a significant decrease in brain area after *CBP3(RNAi)* and *CoREST(RNAi)*. *p≤0.05, ***p≤0.0001 (one-way ANOVA with multiple comparisons). Mean with standard deviation is graphed. Scale bar: 500 µm (A), 100 µm (B), 200 µm (C).

**Figure 3.**
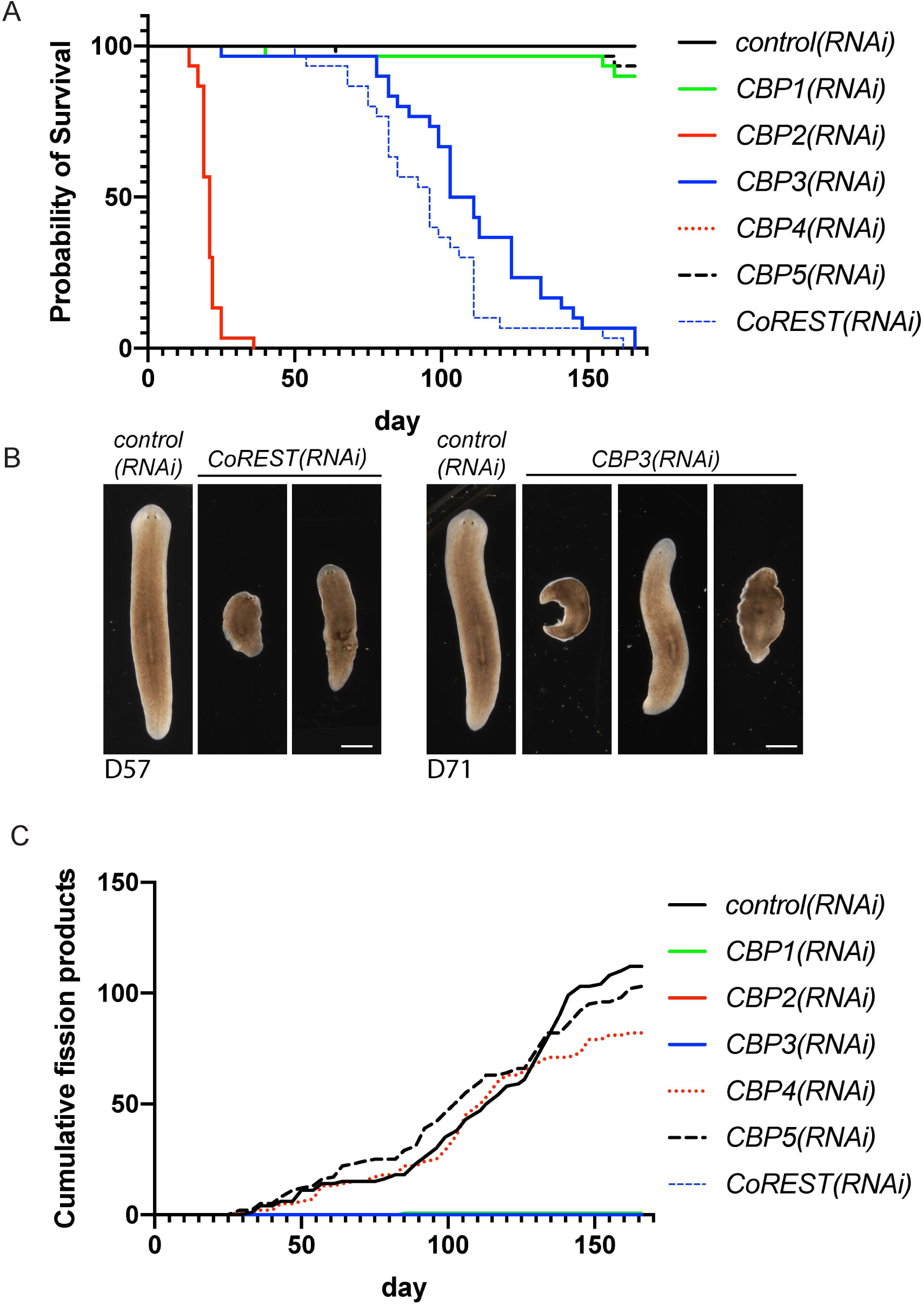
*CBP2, CBP3*, and *CoREST* are important for long-term survival of planarians. (A) A survival curve depicting the relative percentage of surviving animals after long-term RNAi. 30 animals were fed dsRNA once per week for a period of 166 days. Living animals were counted at least twice per week. Death by lysis was fully penetrant for *CBP2(RNAi), CBP3(RNAi)*, and *CoREST(RNAi)* with different rates of lethality. (B) Phenotype analysis of RNAi animals from (A). Head regression was observed in *CoREST(RNAi*) animals at day 57 (D57). A variety of phenotypes including blistering, bloating, and loss of eyespots was observed in *CBP3(RNAi)* animals at day 71. (C) During the survival experiment in (A), fission pieces were counted and discarded at each time point. *CBP4(RNAi)* and *CBP5(RNAi)* animals fissioned at a rate similar to controls. RNAi of *CBP1, CBP2, CBP3*, or *CoREST* inhibited fissioning. Scale bar: 1 mm (B).

Live images were obtained using a Leica DFC420 camera mounted on a Leica M205A stereomicroscope or with an Axiocam color camera mounted on a Zeiss Axio Zoom.V16 microscope. Live video was captured on an iPhone XS and processed in iMovie.

### Immunofluorescence

Planarians were killed in 2% HCl for 5 minutes with alternating 1-minute incubations on ice and shaking at room temperature. Animals were then fixed in 4% formaldehyde solution (in Phosphate Buffered Saline (PBS): 137 mM NaCl, 2.7 mM KCl, 10 mM Na_2_HPO_4_, 2 mM KH_2_PO_4_, pH 7.4) for immunostaining. Animals were bleached under light overnight in 6% H_2_O_2_ in PBS. Bleaching was followed by 3 PBSTx (PBS+0.3% Triton X-100) washes. Animals were then blocked (1X PBS, 0.3% Triton X-100, 0.3g/10ml BSA [Jackson], 0.45% fish gelatin [Sigma]) for at least 4 hr. Blocking solution was replaced with a solution containing primary antibody anti-synapsin (1:250, 3C11 [Developmental Studies Hybridoma Bank]) to visualize the nervous system, and incubated gently shaking at 4°C overnight. Following eight 30-minute PBSTx washes (shaking), animals were incubated in blocking solution for one hour. Blocking solution was replaced with a solution containing the secondary antibody, goat-anti-mouse 488 (1:800, [Jackson]) and animals were allowed to shake gently at 4°C overnight. Animals were washed eight times in PBSTx, for 30-minutes each wash while shaking. In the second wash, DAPI (1 μg/ml) was added to stain cell nuclei. 80% glycerol was used to mount animals for imaging. Images were obtained using Zen software (Carl Zeiss) on a Zeiss LSM 710 confocal microscope with a 20X objective. Images were then processed using ImageJ (Freedman et al., 2003).

### *In situ* hybridization

All riboprobes were generated from PCR products amplified from pJC53.2 clones with T7 primer (see Supplementary Tables 1 and 2). Probes were synthesized with digoxigenin-11-UTP (Roche) at 37°C for 4-5 hr or overnight at 30°C. Probes were treated with RNase-free DNase (Promega) and were cleaned by ammonium acetate precipitation. ISH experiments on whole asexual animals were performed as previously described (King and Newmark, 2013). Briefly, probes were detected with anti-digoxigenin Fab fragments conjugated with alkaline-phosphatase (1:2000, [Roche]). 5-Bromo-4-chloro-3-indolyl phosphate (BCIP, [Roche]) and nitro blue tetrazolium chloride (NBT, [Roche]) in alkaline phosphatase (AP) buffer were used for signal development. Animals were mounted in 80% glycerol and imaged with a Leica DFC420 camera mounted on a Leica M205A stereomicroscope or with an Axiocam color camera mounted on a Zeiss Axio Zoom.V16 microscope. The area of the brain (Fig. 2C-E) or eye (Fig. 6C) was determined after imaging by tracing structures in ImageJ (Schneider et al., 2012) as per (Roberts-Galbraith et al., 2016). *npp-18*^*+*^ and *follistatin*^*+*^ cells were counted for 10 separate 200 µm x 200 µm areas per animal and averaged separately. Average counts were determined for 5 prepharyngeal areas (anterior to the pharynx) and 5 postpharyngeal areas (posterior to the pharynx) for each animal. All statistical analysis on size or cell numbers was conducted through GraphPad Prism software.

### Irradiation

All irradiated animals were exposed to 60 Gray of gamma irradiation, at 150 kV, 5mA on the top shelf of a Gammacell 220 Excel with a cobalt-60 source (Nordion, Ottawa, ON, Canada). Irradiated animals were kept in salts with 1:1000 gentamicin sulfate (50mg/mL stock, [Gemini Bio-Products]) and were later processed for RNA extraction using Trizol Reagent (Invitrogen) either 1 day (∼26 hr) or 5 days (120 hr) post-irradiation.

### Real-time quantitative PCR (RT-qPCR)

Wild type or lethally irradiated asexual animals were used as starting material for isolation of total RNA with Trizol Reagent (Invitrogen) as per the manufacturer’s protocol. An iScript cDNA Synthesis Kit (Bio-Rad) was used to reverse transcribe cDNA from 1µg of RNA using the manufacturer’s protocol. RT-qPCR was performed on an Applied Biosystems StepOne Plus Real-Time PCR system using GoTaq qPCR Master Mix with SYBR Green (Promega) and the primers listed in Supplemental Table 3. All measurements were performed in technical triplicate. Quantities of all transcripts were normalized to the amount of *β-tubulin* mRNA in each sample. Mean values were compared using a Student’s *t* test or one-way ANOVA, as appropriate. More details on statistical analyses are presented in figure legends. All primers used for RT-qPCR are shown in Supp. Table 3.

## Results and Discussion

### Chromatin modifying enzymes function in planarian homeostasis and regeneration

To identify potential epigenetic regulators in planarian tissue homeostasis and regeneration, we first performed a targeted RNAi screen. We first identified a group of 26 planarian homologs of genes that: i) have ties to chromatin modification or remodeling, and ii) are associated with development or disease in other animals (Ballas et al., 2001; Coe et al., 2019; Froyen et al., 2008; Hayami et al., 2011; Husmann and Gozani, 2019; Kraft et al., 2011; Laumonnier et al., 2005; Menke et al., 2016; Montgomery et al., 2009; Petrij et al., 1995; Picketts et al., 1996; Rubinstein and Taybi, 1963; Shi, 2007; Timmermann et al., 2001; Tyagi et al., 2016; Voss and Thomas, 2009; Weiss et al., 2016; Wiszniak et al., 2013; Yu et al., 2010) (Fig. 1A). We cloned each of these genes and examined their functions using RNAi. Animals were fed with dsRNA on days 0, 6, and 12, and homeostasis phenotypes were scored daily. Lysis was detected in *CBP2(RNAi), LSD1b(RNAi)*, and *tip60*(RNAi) animals prior to scheduled amputation (Fig. 1C). We also noted a curling phenotype in *LSD1b*(RNAi) animals (Fig. 1C). For RNAi treatments that resulted in tissue homeostasis problems like blistering or lysis—*ESET(RNAi), LSD1b(RNAi)*, and *tip60(RNAi)*, surviving animals experienced defects in head regeneration including small blastemas after amputation (Fig. 1D, Supp. Fig. 1A). We concluded that several candidates that cause lysis prior to or immediately after amputation—*CBP2, ESET, LSD1b*, and *tip60*—are important for tissue homeostasis in planarians. We note that regenerative phenotypes for these genes may be secondary to tissue damage at the time of amputation.

Because we were also interested in genes that affect planarian regeneration without early tissue damage, we next investigated phenotypes that arise after amputation, specifically focusing on the CBP/p300 family members *CBP1, CBP3, CBP4*, and *CBP5* and the conserved gene *CoREST*, which encodes a histone deacetylase complex member in other species (Andres et al., 1999; You et al., 2001). First, we observed smaller blastema size in *CBP3(RNAi)* animals and *CoREST(RNAi)* animals (Fig. 2A, Supp. Fig. 2A). Next, we wanted to determine whether the regeneration of specific features was perturbed by these gene knockdowns. We examined neural regeneration by amputating RNAi-treated animals and then staining to observe the nervous system using anti-synapsin immunofluorescence (Fig. 2B) and ISH using a *choline acetyltransferase* (*ChAT*) riboprobe (Fig. 2C-E). We noted a strong reduction in anti-synapsin staining in regenerated neural tissue after *CBP3(RNAi)* (Fig. 2B). We also observed a significant reduction in brain area after either *CBP3(RNAi)* or *CoREST(RNAi)* (Fig. 2C). We conclude that perturbation of either *CBP3* or *CoREST* reduces neural regeneration.

Next, to determine whether the functions of CBP genes and CoREST were truly specific to regeneration, we performed long-term knockdown experiments. In these experiments, dsRNA was fed to animals weekly and phenotypes were scored twice weekly. First, we observed animal survival over the study period. Though the timing was somewhat slower than in our previous experiment, we again observed lysis and lethality in *CBP2(RNAi)* animals (Fig 3A). We also saw lethality in *CBP3(RNAi)* and *CoREST(RNAi)* animals, though the timing of death in these treatment conditions was much slower (Fig. 3A). *CoREST(RNAi)* animals showed head regression and smaller size before lysis (Fig. 3B). *CBP3(RNAi)* animals showed a range of phenotypes before lysis, including blistering of the epidermis, loss of one or both eyespots, reduced movement (Supp. Video 1), reduced feeding, and bloating (Fig. 3B).

Due to the duration of the long-term time course, we also observed fissioning of control animals over time. As fission pieces were formed, they were counted and removed from the Petri dishes twice per week. We observed that *CBP3(RNAi)* and *CoREST(RNAi)* animals never fissioned, consistent with their lethal phenotypes and shrinking size over time. *CBP4(RNAi)* and *CBP5(RNAi)* animals fissioned frequently over the time course, similar to control animals (Fig. 3C). Surprisingly, though *CBP1(RNAi)* animals showed no overt phenotypes, they failed to produce fission pieces (only 1 fission event during the entire time course vs. >100 for the control(RNAi) population, Fig. 3C). *CBP1(RNAi)* animals maintained a similar size over the course of the experiment, suggesting that more subtle defects in feeding, growth, or asexual reproduction might occur in the absence of the *CBP1* gene product.

Taken together, our results indicate *CoREST* and the diversified *CBP* family in planarians play novel roles in regeneration, growth, and homeostasis in planarians. In our primary screen, we also confirmed phenotypes of six other genes — listed in blue — that have been reported elsewhere (*HDAC1, CHD4, trr-1, MLL1/2, SET1*, and *HUWE1*) (Henderson et al., 2015; Hubert et al., 2013; Scimone et al., 2010; Zhu and Pearson, 2013). The remaining fourteen genes identified in this screen and listed in black (*CBP5, CBP4, MYST4, GCN5, Jarid1, Jarid2, PHF8, ATRX1, CHD7, SNF2, SMARCAL1*, and *NEDD4L*) or previously published and represented in grey (*trr-2*) (Hubert et al., 2013), produced either no or very subtle phenotypes in our RNAi experiments.

### Expression patterns of putative chromatin modifiers in planarians

We were interested in understanding further the potential roles of the expanded CBP/p300 family in planarians. First, to assess whether the CBP genes were predominantly expressed in stem cells, we determined their expression patterns using whole mount ISH (Fig. 4A). The five planarian *CBP* genes were expressed broadly throughout the animal, with *CBP1, CBP2, CBP3*, and *CBP4* enriched in the cephalic ganglia and *CBP5* enriched in the parenchyma, the location of planarian stem cells (Fig. 4A). The broad expression patterns of the *CBP* genes suggest that these histone acetyltransferases could function in multiple cell types.

**Figure 4.**
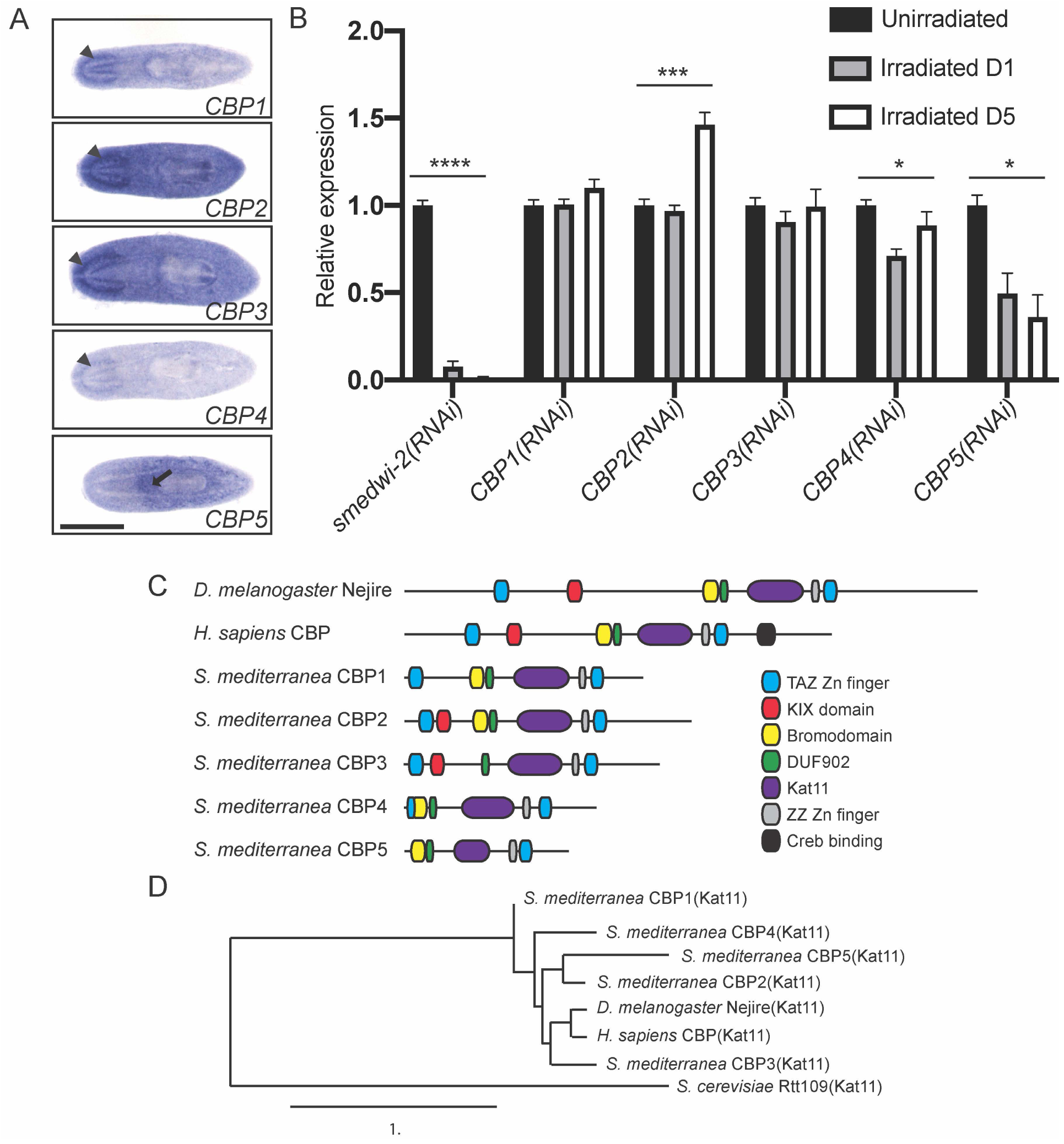
The *CBP/p300* family is expanded in planarians. (A) ISH showing expression patterns for the 5 planarian *CBP* genes. The *CBP* genes are broadly expressed throughout the animal. In addition to this broad expression pattern, *CBP1, CBP2, CBP3*, and *CBP4* are enriched in the brain (arrowheads) while *CBP5* is enriched in the parenchyma (arrow). (B) Relative expression levels of *CBP* genes were measured by RT-qPCR performed on cDNA preparations from unirradiated animals or animals either 1 or 5 days after irradiation for ablation of stem cells. A significant decrease in *smedwi-1* and *CBP5* mRNA was observed, with no steady decrease for *CBP1, CBP2, CBP3*, and *CBP4* transcripts. Measurements were performed in triplicate (n=10 animals each). Error bars are SEM. *p≤0.05, ***p≤0.001, ****p≤0.0001 (one-way ANOVA). (C) The domain architecture of CBP family proteins is depicted. Domains were identified with Pfam and were compared between *Drosophila melanogaster, Homo sapiens*, and *Schmidtea mediterranea* CBP orthologs. (D) Phylogenetic tree of CBP family orthologs. Analysis was based on the chromatin-acetylating Kat11 domain. The Kat11 domain of *S. cerevisiae* Rtt109 was used as an out-group. Scale bar (A): 500 µm.

We further sought to confirm whether *CBP5* is enriched in the stem cells, which can be ablated by irradiation. Following irradiation, RT-qPCR analysis showed a significant decrease in *smedwi-1* mRNA levels (Fig. 4B). Since *smedwi-1* mRNA is expressed in stem cells, reduction in *smedwi-1* transcript levels after irradiation indicates stem cell ablation (Reddien et al., 2005b). mRNA levels remained mostly constant or slightly increased for *CBP1, CBP2, CBP3*, and *CBP4* after irradiation, consistent with their broad expression by ISH. The exception was an observed decrease in *CBP5* mRNA levels after stem cell ablation. When combined with the ISH results, the data suggest that *CBP5* is enriched in stem cells. These results are also corroborated by previously published sequencing analyses performed on sorted stem cells and single cells, which indicate that all five *CBP* paralogs are expressed in stem cells but also in many other cell types (Supp. Table 1; (Fincher et al., 2018; Labbé et al., 2012)).

In addition to assessing the expression patterns of *CBP* orthologs, we also determined the expression patterns of the other novel genes with roles in planarian homeostasis and regeneration. We saw that *ESET, LSD1b*, and *tip60* are broadly expressed in the planarian, while *CoREST* is more specifically enriched in the nervous system (Supp. Fig. 1B). Using datasets generated to investigate gene expression both at the population and single-cell levels, nearly all the genes in our list were expressed in stem cells and often in other cell types, supporting the idea that chromatin dynamics are occurring during stem cell maintenance and differentiation (Supp. Table 1 (Fincher et al., 2018; Labbé et al., 2012)).

### CBPs and their domains in *S. mediterranea*

In metazoans, CBPs regulate gene expression (Vo and Goodman, 2001) by functioning as scaffolds to bridge diverse transcriptional activators and coactivators (Hecht et al., 2000; Kwok et al., 1994). In addition, CBPs regulate transcriptional coactivators and chromatin itself via their acetyltransferase activity (Bannister and Kouzarides, 1996; Ogryzko et al., 1996; Raisner et al., 2018; Weinert et al., 2018). CBPs possess multiple distinct functional domains, which allow them to interact with many different proteins simultaneously to perform these functions.

The human genome encodes two CBP-like genes, *CBP* and *p300*. Germline mutations in human *CBP* result in Rubinstein–Taybi syndrome, characterized by intellectual disability and predisposition to malignancies in early life (Petrij et al., 1995). Mice also have two CBP-like genes (also *CBP* and *p300*), with loss of both *CBP* alleles or compound heterozygosity causing fatality (Yao et al., 1998). *Caenorhabditis elegans* c*bp-1*, the single *CBP/p300* homolog in this species, reverses the repressive effects of histone deacetylases to promote transcriptional activation and differentiation during early embryogenesis (Shi and Mello, 1998) while the *Drosophila* homolog *Nejire* is a co-activator of Cubitus interruptus in the Hedgehog signaling pathway and is important for pattern formation (Akimaru et al., 1997a; Akimaru et al., 1997b). In parasitic flatworms, a CBP homolog, *CBP1*, was found to regulate schistosome stem cell function (Bertin et al., 2006; Collins and Collins, 2016). Since CBP proteins function with a variety of partners in diverse biological processes, *cbp* mutants are often pleiotropic (Janknecht and Hunter, 1996). In contrast, the planarian genome encodes <u>five</u> paralogous *CBP* genes. We reasoned that the five *CBP* paralogs might offer an *in vivo* system in which the functions of the CBP genes may be more clearly assessed.

To begin characterizing the planarian CBP family, we identified the domain architecture for each paralog and compared them to *Drosophila* and human homologs (Fig. 4C). All CBP proteins contain the Kat11 domain, which is the catalytic domain responsible for acetyltransferase activity. In other organisms, this domain functions to acetylate residues on histones and non-histone proteins (e.g. K56 on histone H3), often to promote transcriptional activation (Das et al., 2009; Raisner et al., 2018). Another domain shared by all orthologs is the TAZ (Transcription Adapter putative Zinc Finger) domain. This domain is present on the N- and C-terminal ends of each ortholog except *S. mediterranea* CBP5, and forms a structure that mediates protein-protein interactions for mammalian CBP and p300 (De Guzman et al., 2005a; De Guzman et al., 2005b). Yet another shared domain is the ZZ-type Zn finger, which is thought to be involved in protein-protein interactions as well (Legge et al., 2004; Zhang et al., 2018).

Despite the similarities, certain domains are encoded by some but not all orthologs. The KIX domain, found in *D. melanogaster* Nejire, *H. sapiens* CBP, and *S. mediterranea* CBP2 and CBP3, mediates interactions between characterized CBPs and transcription factors like c-Myb and CREB (cAMP response element binding protein; (Radhakrishnan et al., 1997; Toto et al., 2014)). Bromodomains are present in all orthologs except *S. mediterranea* CBP3. Bromodomains mediate binding previously acetylated lysine residues on histone tails or on transcription factors (Dhalluin et al., 1999; Polesskaya et al., 2001; Zeng et al., 2008); the presence of a bromodomain in planarian CBP orthologs may confer affinity to acetylated chromatin regions or acetylated transcription factors to propagate acetylation regionally. We also observed that the Kat11 domain is truncated in CBP5, which could indicate a difference in catalytic activity.

To further assess the relationships between *D. melanogaster, H. sapiens*, and *S. mediterranea* homologs, we performed a phylogenetic analysis (Fig. 4D). The Kat11 domain was used for comparison with *S. cerevisiae* Rtt109(Kat11) as an out-group. *D. melanogaster* CBP(Kat11) and *H. sapiens* CBP(Kat11) were most closely related to each other, and then to *S. mediterranea* CBP3(Kat11). Among the *S. mediterranea* paralogs, we observed the closest relationship between *S. mediterranea* CBP2(Kat11) and CBP5(Kat11). Together, the variation in domain architecture and the evolutionary relationship between the CBPs in *Drosophila melanogaster, Homo sapiens*, and *Schmidtea mediterranea* support a diversified CBP family in planarians.

### *CBP2* and *CBP3* are necessary for distinct stem cell functions

The differences we detected between *CBP2* and *CBP3* in their domain architecture (Fig. 4C) and RNAi phenotypes (Fig. 1-3) led us to hypothesize that the two genes have evolved separate functions in planarians. Due to the quick and severe homeostatic defects detected in *CBP2(RNAi)* animals, we first examined whether *CBP2* affected stem cell maintenance or function. Indeed, we observed a striking decrease in the stem cell marker *smedwi-1* after *CBP2(RNAi)* (Fig. 5A), suggesting that CBP2 affects stem cell survival. We confirmed the loss of stem cells after *CBP2(RNAi)* by performing RT-qPCR to detect a variety of stem cell-enriched transcripts (Fig. 5B (Reddien et al., 2005b; van Wolfswinkel et al., 2014; Wagner et al., 2012)). We saw a decrease in both broadly expressed stem cell markers (*smedwi-1, smedwi-2, H2B*) and markers like *zfp-1* that is enriched more specifically in the epidermal lineage of stem cells in the planarian (Fig. 5B (van Wolfswinkel et al., 2014)). Other stem cell marker genes (transcription factor-encoding genes *soxP-1, soxP-2*; and FGF-receptor encoding genes *fgfr-1* and *fgfr-2*) were all affected, though to varying degrees (Fig. 5B). We did not observe a decrease in markers of differentiated cells like *choline acetyltransferase* (*ChAT*, Fig. 5B). We thus conclude that *CBP2* is essential for stem cell maintenance in planarians.

**Figure 5.**
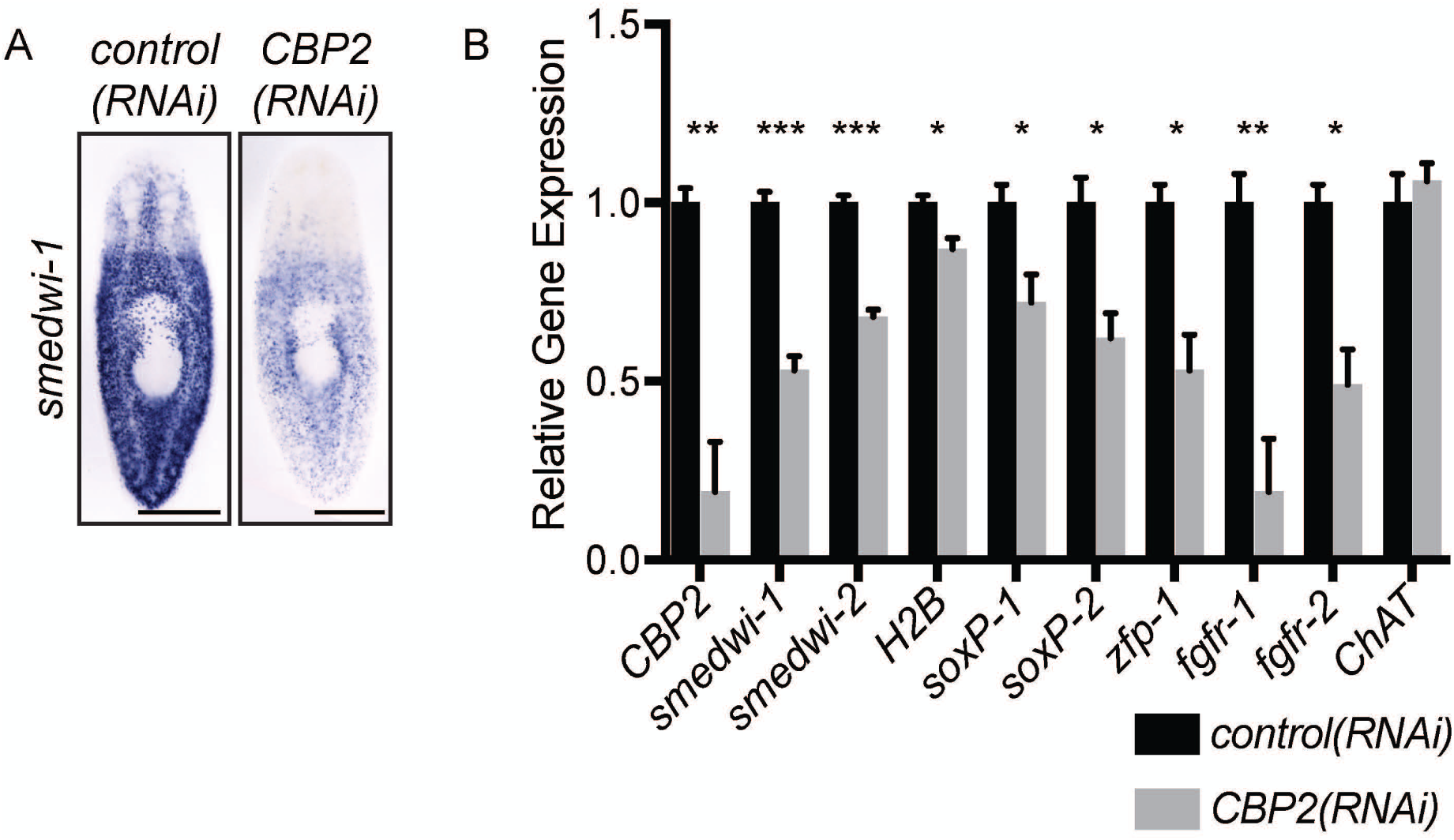
Knockdown of individual *CBP/p300* orthologs causes defects in planarian homeostasis and regeneration. (A) ISH was used to visualize the stem cell marker *smedwi-1* in non-amputated *control(RNAi)* and *CBP2(RNAi)* animals. (B) RT-qPCR was used to detect levels of *CBP2* transcript, a variety of stem cell-enriched transcripts after RNAi. The neural marker *choline acetyltransferase* (*ChAT*) was used as a non-stem cell control. Measurements were performed in biological and technical triplicates. Error bars are SEM. *p≤0.05, **p≤0.01, ***p≤0.001 (student t-tests). Scale bar: 500 µm (A).

We next turned our attention to *CBP3*, which we had already shown affected brain regeneration (Fig. 2) and long-term survival of planarians (Fig. 3). We first examined whether stem cell maintenance was affected in *CBP3(RNAi)* animals as it is after *CBP2(RNAi)*. While *CBP3(RNAi)* caused a reduction in stem cell marker *smedwi-1* by ISH and RT-qPCR (Fig. 6A-B), the reduction was milder than that seen after *CBP2(RNAi)* (Fig. 5). We next examined whether stem cell function might be more narrowly affected after *CBP3(RNAi)*. Using RT-qPCR, we found that several genes expressed in neurons and neural progenitors (*ap-2, klf, lhx1/5-1, soxB2-2;* (Currie and Pearson, 2013; Roberts-Galbraith et al., 2016; Scimone et al., 2014; Wenemoser et al., 2012)) and eyespots and eye progenitors (*ovo, klf; (Lapan and Reddien, 2012)*) were significantly downregulated after *CBP3(RNAi)* (Fig. 6B). We confirmed the decrease in eye markers by ISH and observed that *CBP3(RNAi)* resulted in smaller eyespots in regenerating animals (Fig. 6C). We sought to confirm the reduction in neural progenitor markers and did see a reduction in *ap-2* staining through ISH (Fig. 6D); *ap-2* is expressed in both neural progenitors and some mature neural cell types and *ap-2* is required for regeneration of *TrpA*^*+*^ neurons (Scimone et al., 2014; Wenemoser et al., 2012). Further, *neuroD-1*, a gene expressed in neurons and putative neural progenitors (Fig. 6E (Scimone et al., 2014)), was previously reported to impact neural specification (Cowles et al., 2013). In more recent single-cell sequencing analysis, *neuroD-1, neuropeptide precursor-18 (npp-18)*, and *follistatin* were all shown to be co-expressed in a subcluster of neurons (subcluster 28, (Fincher et al., 2018)). Based on our prediction that *neuroD-1* might promote formation or maintenance of this cell type, we knocked down *neuroD-1* and showed that RNAi indeed affects *npp-18*^+^ cells (Fig. 6F-I) and *follistatin*^*+*^ cells (Supp. Fig. 3), in both new and old tissue. We therefore used *neuroD-1* as an additional marker of neural lineages and found that *CBP3(RNAi)* also strongly reduced *neuroD-1* (Fig. 6J).

**Figure 6.**
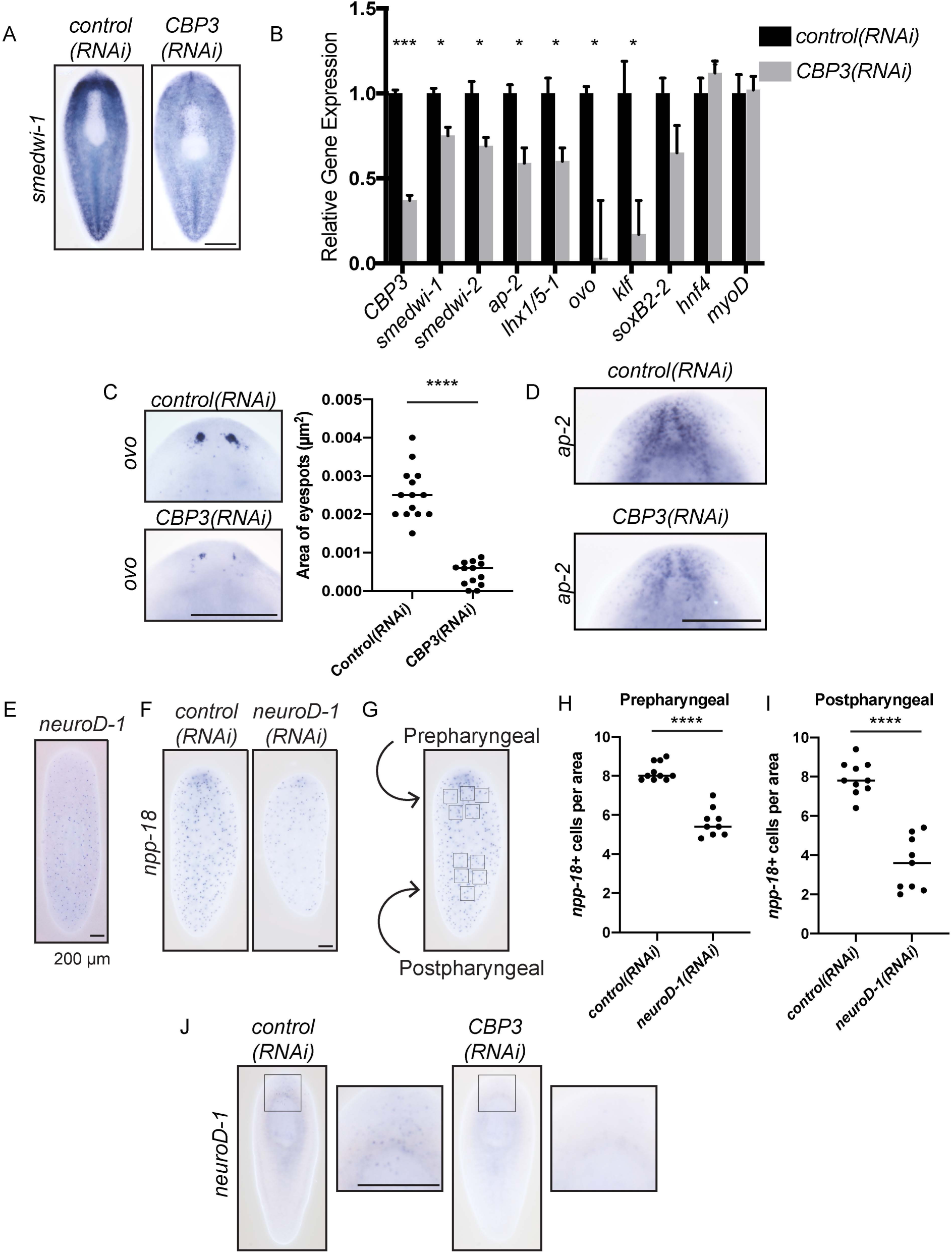
*CBP3* plays a restricted role in stem cell progeny. A) *control(RNAi)* and *CBP3(RNAi)* animals were stained using ISH with the stem cell marker *smedwi-1. smedwi-1* staining is somewhat decreased in *CBP3(RNAi)* animals 7 dpa. (B) *control(RNAi)* and *CBP3(RNAi)* animals were fed dsRNA three times over 10 days, amputated on day 16, and processed on day 5 after amputation for RNA purification and RT-qPCR. Measurements were performed in biological and technical triplicates. Error bars are SEM. We observed significant decreases in stem cell markers *smedwi-1* and *smedwi-2*. We observed decreases in *ap-2, klf, lhx1/5-1*, and *soxB2-2*, which each mark subsets of neurons and neural progenitors. We also observed decreases in *ovo* and *klf* which mark eye cells and progenitors. We did not see a difference in *hnf4*, which marks intestinal progenitors or *myoD*, which marks muscle progenitors. *p≤0.05, ***p≤0.001 (multiple *t*-tests). C) *ovo* is used to mark eyespot cells and progenitor trails in regenerating animals 7 dpa. The area of *ovo*-marked eyespots were quantified after *control(RNAi)* and *CBP3(RNAi)* and was normalized against body size. Eyespots were significantly smaller after regeneration in *CBP3(RNAi)* animals (p≤0.0001; n=13-14; Mann-Whitney test). (D) *control(RNAi)* and *CBP3(RNAi)* animals were stained via ISH with *ap-2*, a marker of neurons and neural progenitors. A decrease in *ap-2* signal was seen after *CBP3(RNAi)*. (E) The expression pattern of *neuroD-1* is shown in non-amputated animals by ISH. (F) *control(RNAi)* and *neuroD-1(RNAi)* animals were subjected to ISH using an *npp-18* riboprobe. Fainter *npp-18* staining is evident after *neuroD-1(RNAi)*. (G) Illustration showing how 5 200 µm x 200 µm boxes were drawn in both prepharyngeal and postpharyngeal areas of each animal to quantify cell number. *npp-18*^*+*^ cells were counted in each area and averaged for each animal (n=10-11 animals per condition). (H-I) *neuroD-1(RNAi)* animals had fewer *npp-18*^*+*^ cells in both prepharyngeal and postpharyngeal regions. (****, P<0.0001, Welch’s t-test). (J) ISH was performed to detect *neuroD-1* at 7 dpa after *control(RNAi)* or *CBP3(RNAi)* treatment at 7 dpa. *neuroD-1*^+^ cells are markedly decreased after *CBP3(RNAi)*. (H Scale bars: 500 µm (A, C, D, J); 200 µm (E, F).

Taken together, we observed that CBP3 affects brain regeneration (Fig. 2) and is important for proper expression of several neural and neural progenitor markers (Fig. 6). We reasoned that CBP3 might affect neural lineages specifically or might affect stem cell differentiation more broadly. We next examined other markers of non-neural cell lineages in planarians. We did see a decrease in one early epidermal lineage marker after *CBP3(RNAi)* (*prog-1*, Supp. Fig. 2C (Eisenhoffer et al., 2008)). This observation is consistent with lysis we observed after long-term RNAi (Fig. A-B). However, we also determined that markers of gut progenitors (*hnf4*; (Scimone et al., 2014)), muscle progenitors (*myoD;* (Scimone et al., 2017; Scimone et al., 2014)), and epidermal progenitors (*agat-3*; (Eisenhoffer et al., 2008)) appeared unaffected after *CBP3(RNAi)* (Fig. 6B, Supp. Fig. 2D). Therefore, we conclude that *CBP3* affects stem cell maintenance and differentiation with a narrow influence on neuronal and potentially epidermal lineages.

## Conclusion

In our search for novel chromatin modulators, we identified several novel genes important for aspects of planarian physiology. Importantly, our data indicate that the CBP family of histone acetyltransferases has diversified in planarians. Our findings support distinct roles for CBP2 and CBP3 in stem cell maintenance and differentiation. Our data argue that CBP3 functions more narrowly, preferentially affecting neural fates, whereas CBP2 plays an essential role in stem cell maintenance. Finally, though we did not focus on the characterization of the other CBP genes (*CBP1, CBP4*, and *CBP5*), the fission defect after *CBP1(RNAi)* could suggest that additional planarian CBP proteins might have unidentified functions. Previous work suggests that homologous proteins in other organisms, such as the *Drosophila* Nejire and the *C. elegans* Cbp-1, function in differentiation pathways during embryonic development (Akimaru et al., 1997a; Akimaru et al., 1997b; Shi and Mello, 1998), suggesting potentially conserved roles for this gene family. Pleiotropic consequences are observed in other model organisms upon knockdown of *CBP* genes, thus investigating the division of labor between *CBP*s in planarians could uncover conserved roles in stem cell differentiation in other systems. In the future, it will be important to identify molecular roles of CBP2 and CBP3—including verifying their roles in chromatin modification and transcriptional regulation in planarian cells—and to characterize the properties that confer these factors with distinct functions in stem cell biology.

---

**Supplemental Figure 1.**
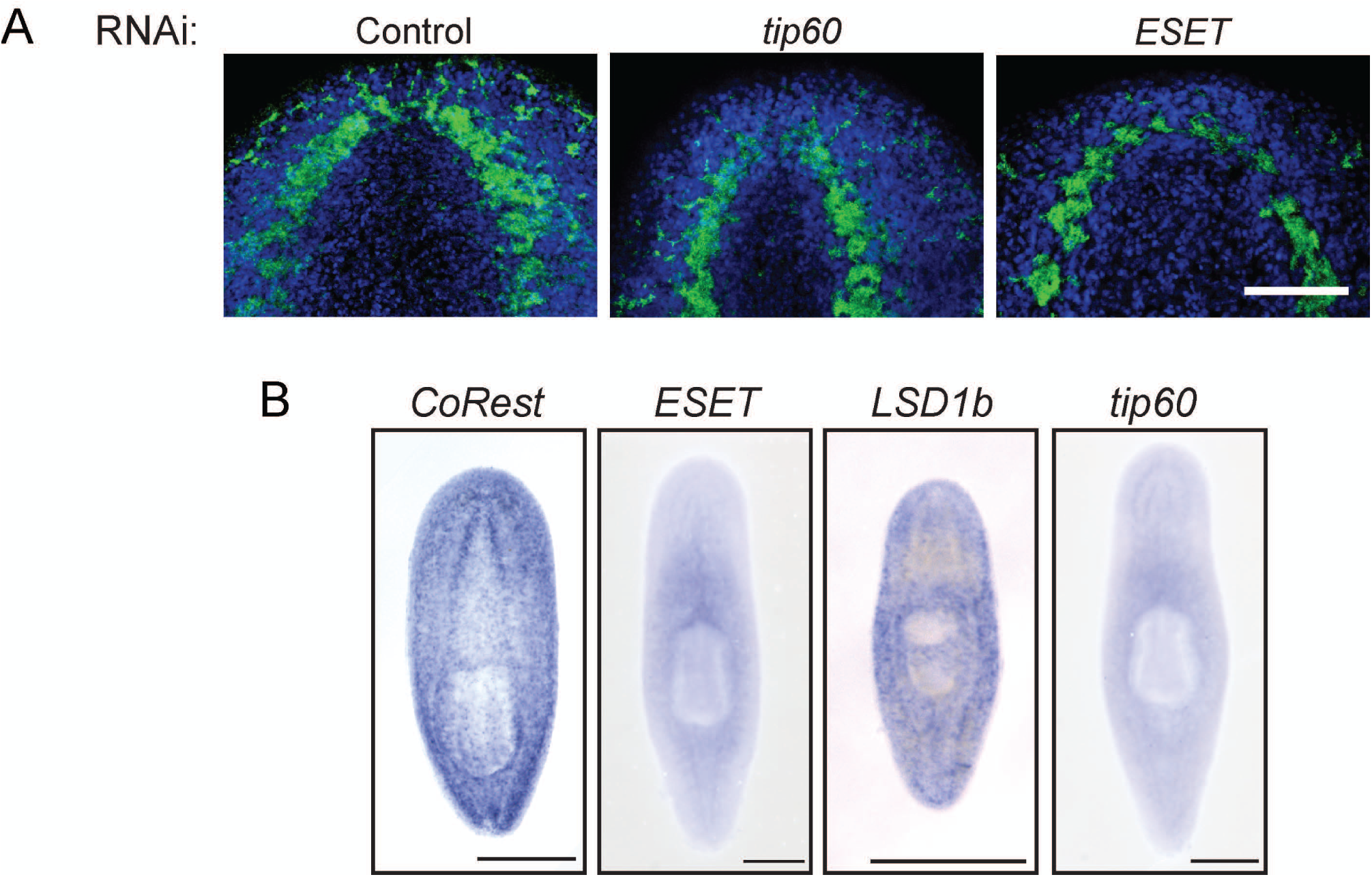
Chromatin modifying and remodeling enzymes are important in planarian homeostasis and regeneration. A) Animals fed control dsRNA or dsRNA matching *tip60* or *ESET* (as per the paradigm in Fig. 1B) were amputated and allowed to undergo head regeneration. Regenerates were stained with anti-synapsin (green) and DAPI (blue). (B) ISH showing the expression pattern for the indicated genes. The genes are broadly expressed with enrichment in the brain for *CoREST*. Scale bar: 100 µm (A), 500 µm (B).

**Supplementary Figure 2.**
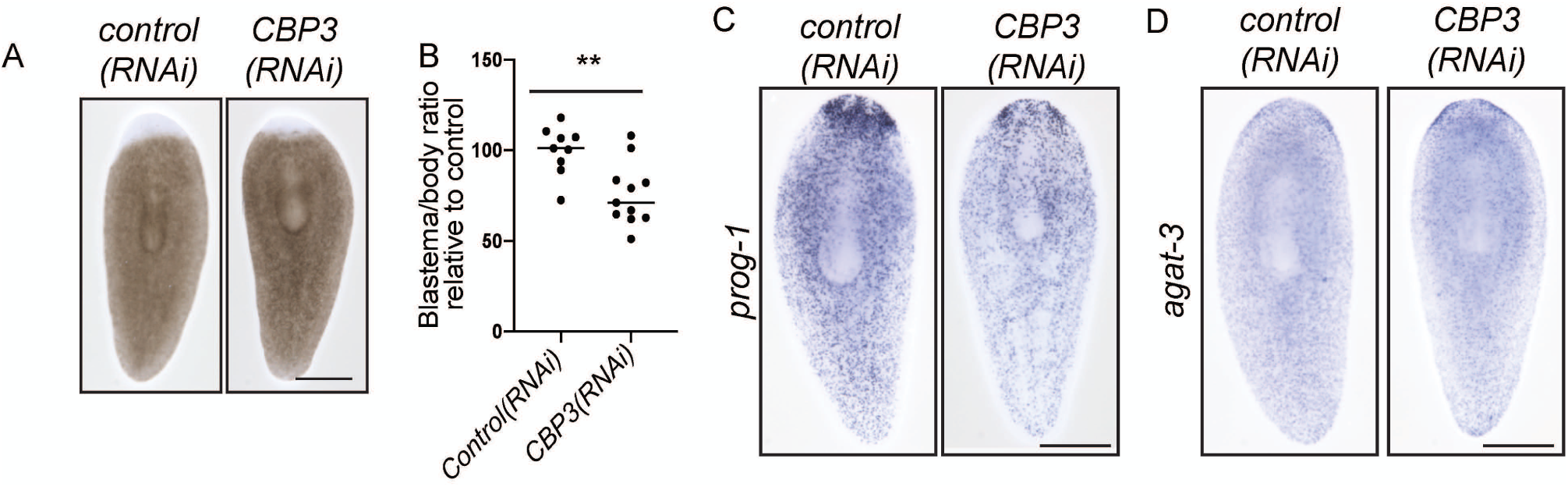
CBP3 has a narrower role in regeneration. (A) Blastemas were imaged in fixed *control(RNAi)* and *CBP3(RNAi)* animals. (B) Blastemas from (A) were measured and calculated as a percentage of animal body size. We graphed the results normalized so that the average blastema size in control animals appears as 100%. Blastema size was significantly smaller after *CBP3(RNAi)*. (n=9, 11; ** indicates P=0.0030 using a student’s t-test.) (C-D) Epidermal lineage markers *prog-1* and *AGAT-3* were visualized by ISH after *control(RNAi)* and *CBP3(RNAi)* in regenerating worms 7 dpa. Scale bars: 500 µm.

**Supplemental Figure 3.**
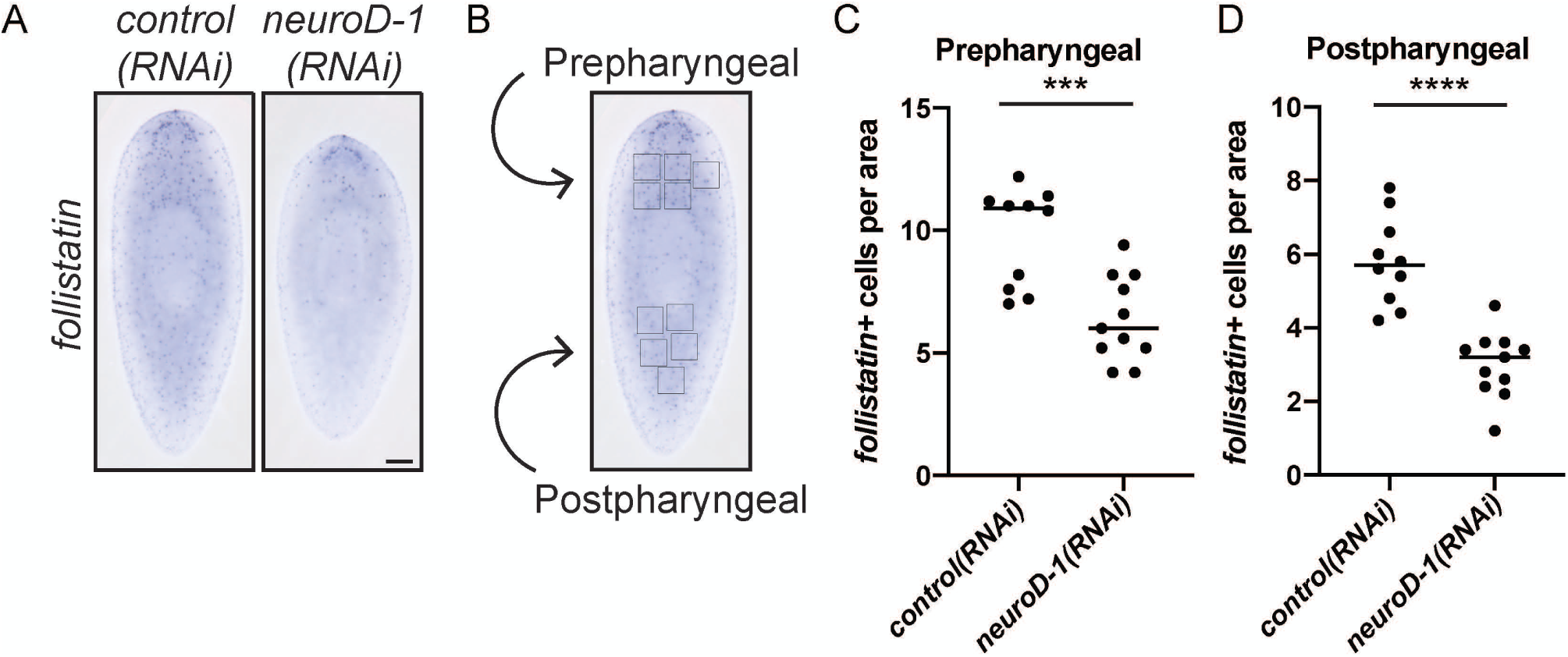
NeuroD-1 influences medial follistatin^+^ cells in the planarian body. (A) *control(RNAi)* and *neuroD-1(RNAi)* animals were subjected to ISH using a *follistatin* riboprobe. *follistatin* staining at the anterior pole is similar after *neuroD-1(RNAi)*, but fewer *follistatin*^*+*^ cells in the posterior are visible. (B) Illustration showing how 5 separate 200 µm x 200 µm boxes were drawn in each prepharyngeal and postpharyngeal areas of each animal to quantify cell number. *follistatin*^*+*^ cells were counted in each area and averaged for each animal (n=9-10 animals per condition). (C-D) *neuroD-1(RNAi)* animals had fewer medial *follistatin*^*+*^ cells in both prepharyngeal and postpharyngeal regions. (***, P<0.001; ****, P<0.0001, Welch’s t-test).

Supplementary Table 1. ***Genes from this study***. This table includes gene identifications, primer sequences for cloning, and homology details for all genes from this study.

Supplementary Table 2. ***Primers used for riboprobe and dsRNA synthesis in this study***.

Supplementary Table 3. ***Primers used for RT-qPCR in this study***.

Supplementary Video 1. ***Size and movement defects in CBP3(RNAi) animals***. A 2-minute video was created of *control(RNAi)* and *CBP3(RNAi)* animals at D64 of the long-term RNAi experiment described in Fig. 3.

## Acknowledgments

The authors thank past members of the Newmark laboratory for their feedback in the early phases of this project. CRS was supported during her studies by the James Scholar Summer Research Award, the Molecular and Cellular Biology Summer Fellowship, the Jenner Family Fellowship, and the Office of Undergraduate Research Award. RRG was supported by start-up funds from the University of Georgia.

